# Functional rescue of a fatal ERAD mutation via alternative splicing

**DOI:** 10.1101/2025.06.13.659581

**Authors:** Huilun Helen Wang, Zhihong Wang, Liangguang Leo Lin, Sunil K Verma, Weronika Gniadzik, Hui Wang, Zexin Jason Li, Lulu Jiang, Muge N Kuyumcu-Martinez, Shengyi Sun, Ling Qi

**Affiliations:** Department of Molecular Physiology and Biological Physics, University of Virginia School of Medicine, Charlottesville, VA 22903, USA; Department of Neuroscience, Center for Brain Immunology and Glia (BIG), University of Virginia School of Medicine, Charlottesville, VA 22908, USA; Robert M. Berne Cardiovascular Research Center, University of Virginia School of Medicine, Charlottesville, VA 22903, USA; Center for RNA Science and Medicine, University of Virginia School of Medicine, Charlottesville, VA 22903, USA; Compherensive Cancer Center, University of Virginia School of Medicine, Charlottesville, VA 22903, USA; Department of Pharmacology, University of Virginia School of Medicine, Charlottesville, VA 22903, USA

**Author notes:** Correspondence (H.H.W.); (S.S.); (L.Q.).

**Keywords:** ER-associated degradation, SEL1L, disease variant, ENDI-A, mouse models, alternative splicing, B cell lymphopenia, neurodevelopment, anti-sense oligonucleotide, patient fibroblasts

## Abstract

Endoplasmic reticulum (ER)-associated degradation (ERAD) is essential for cellular proteostasis, with the SEL1L-HRD1 protein complex targeting misfolded proteins in the ER for proteasomal degradation. Disruption of this pathway underlies a recently identified infant-onset neurodevelopmental disorder (ENDI syndrome), characterized by profound developmental delay, microcephaly, and immune deficiency. Its most severe form, ENDI with agammaglobulinemia (ENDI-A), is driven by a bi-allelic SEL1L Cys141Tyr (C141Y) mutation within the fibronectin II (FNII) domain, for which no treatment currently exists. Here, we serendipitously uncover a striking mechanism of intrinsic rescue in knock-in mouse models of the C141Y mutation: enhanced usage of an alternative splice donor site within exon 4 bypasses the mutant FNII-encoding region, restoring ERAD activity and rescuing key disease phenotypes including perinatal lethality, growth retardation, B cell deficiency, and neurodevelopmental defects. Building on this discovery, we demonstrate that antisense oligonucleotide (ASO)-mediated exon skipping in patient-derived fibroblasts generates a truncated yet functional SEL1L protein, fully rescuing ERAD function and ER proteostasis. These results establish RNA splicing modulation as a viable therapeutic strategy for ERAD deficiency and extend the clinical potential of exon-skipping therapy to diseases of protein misfolding.

**ONE-SENTENCE SUMMARY:** This study reports the discovery of using antisense oligonucleotides (ASOs) to rescue the biallelic SEL1L C141Y variant, offering a potential therapeutic strategy.

## MAIN TEXT

Nascent proteins in the endoplasmic reticulum (ER) that fail to fold properly are recognized as misfolded and subsequently targeted for cytosolic proteasomal degradation by an ER quality control system known as ER-associated degradation (ERAD) (*1–3*). In mammals, the SEL1L-HRD1 protein complex represents the most conserved branch of ERAD (*1, 4, 5*). Misfolded proteins are first recognized by lectins such as OS9 and then delivered to SEL1L, which are then subsequently retrotranslocated through the HRD1 retrotranslocon followed by HRD1-mediated ubiquitination and proteasomal degradation (*6–8*). SEL1L is required for substrate recruitment, HRD1 protein stability (*9, 10*), and ERAD complex formation (*11–13*). Using various *Sel1l-* or *Hrd1-* knockout mouse models, we and others have shown that the SEL1L-HRD1 ERAD pathway plays vital roles in a range of physiological processes in a substrate- and cell-type-specific manner, largely uncoupled from ER stress response (*14–16*).

The recent identification of human patients carrying bi-allelic *SEL1L* and *HRD1* variants has provided direct evidence of its critical role in humans. To date, eleven patients have been found carrying four distinct bi-allelic hypomorphic variants in *SEL1L* and *HRD1*, all presenting with moderate to severe intellectual disability, microcephaly, developmental delay and locomotor dysfunction, collectively termed ENDI (*17, 18*). Among affected individuals, those carrying the SEL1L C141Y mutation (NM_005065.6: exon 4: c.422G>A, p.Cys141Tyr) exhibit not only the core symptoms of ENDI but also agammaglobulinemia and early mortality. This more severe phenotype, termed ENDI-agammaglobulinemia (ENDI-A), has been associated with heightened ERAD dysfunction (*18*). However, despite these insights, direct evidence supporting the pathogenicity of this variant remains limited, and no therapeutic strategies have been established to date.

In this study, while establishing the disease causality of the SEL1L C141Y mutation, we unexpectedly discovered that modulating splicing at exon 4 could reverse disease pathology in a knock-in (KI) mouse model carrying the mutation. Building on this finding, we successfully restored ERAD function in SEL1L C141Y patient-derived human fibroblasts using splice-switching antisense oligonucleotide (ASO)- mediated exon skipping. While modulating alternative splicing using ASOs – short, synthetic nucleotides that bind target RNAs to alter splicing or promote degradation – has emerged as a powerful therapeutic approach for genetic diseases (*19–21*), our findings extend this strategy to target ERAD dysfunction, an area previously unexplored.

### Alternative splicing of *Sel1L* exon 4 rescues lethality in SEL1L C141Y KI mice

C141 is located within the fibronectin II (FNII) domain of SEL1L (**Fig. 1A**), a ∼50-amino acid motif of unknown function that is notably absent in yeast SEL1L ortholog Hrd3 (*18, 22*). This domain, encoded by exon 4 (**Fig. 1A**), is stabilized by two disulfide bonds (Cys127-Cys153 and Cys141-Cys168) (*23*), with C141 forming part of the latter (**fig. S1A**). To investigate the disease causality and pathological consequences of *SEL1L* C141Y mutation (corresponding to C137Y in mice) *in vivo*, we generated KI mouse models using CRISPR-Cas9 genome editing (**fig. S1B**). Sanger sequencing confirmed successful introduction of the intended G-to-A mutation in the genomes of three independent founder lines (Lines A-C) (**Fig. 1B** and **fig. S1C**), from which we obtained WT, heterozygous (HET), and homozygous KI offspring (**fig. S1, D to E**). Unexpectedly, only Line C produced viable homozygous mice (hereafter KI’) at the expected ratio that survived past postnatal day 21 (p21) at Mendelian ratios. In contrast, Lines A and B yielded only 5 viable homozygous mice out of ∼ 280 pups surviving past weaning age p21 – an overall survival rate of ∼ 1.8% (**Fig. 1C**). HET mice from all lines were viable (**Fig. 1C**) and appeared phenotypically indistinguishable from WT littermates. Given their shared phenotypes, they were grouped as the KI line for subsequent analyses using tissues from p0 neonates due to lethality.

**Fig. 1.**
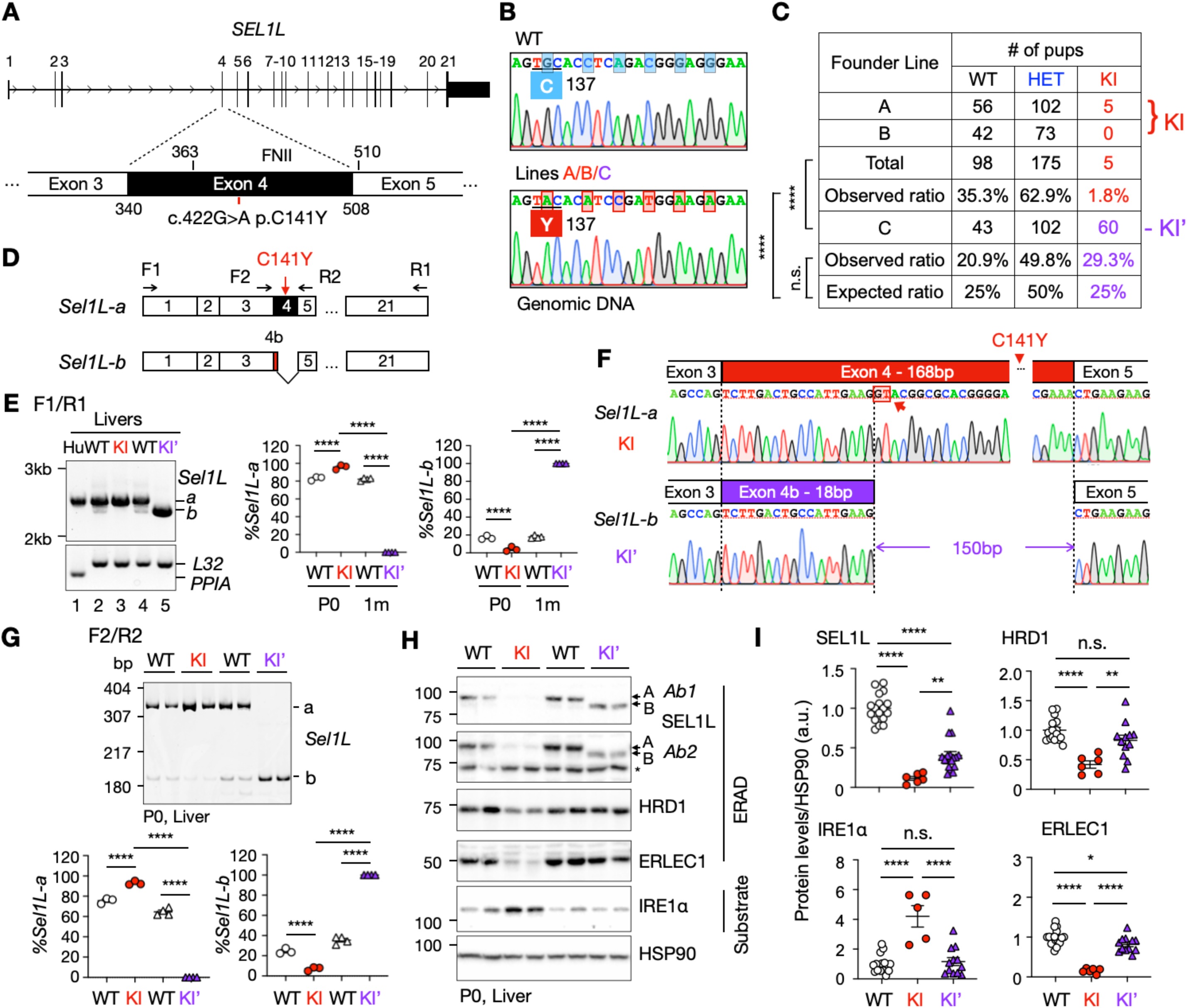
The alternative splicing of Sel1L exon 4 rescues ERAD function and lethality of SEL1L C141Y knockin (KI) mice. **(A)** A diagram showing the location of *SEL1L* C141Y mutation (red line) and Fibronectin II (FNII) domain on SEL1L gene. **(B)** Sanger sequencing of *Sel1L* genes from the homozygous progenies of WT and KI founder lines A/B/C. Red shaded box highlights the mutation identified in exon 4. **(C)** The survival rate of KI/KI’ pups and their WT and HET littermates at postnatal day 21 (P21). **(D)** The diagram of the Sel1L isoforms and the primer design for **(E)** and **(G)**. **(E)** DNA agarose gel electrophoresis analysis and quantitation of full length *SEL1L* transcript from WT human livers and *Sel1L* transcripts from WT, KI and KI’ mouse livers at ages of postnatal day 0 (P0) and 1 month (1m), respectively. *PPIA* and *L32,* internal controls for human and mouse tissues, respectively. n = 3-4 mice/group. **(F)** Sanger sequencing of SEL1L cDNA, focusing on exon 4 and splicing junctions. Red arrow and shaded red box, the alternative GT donor. **(G)** DNA polyacrylamide gel electrophoresis (PAGE) analysis and quantitation of *Sel1L* exon 4 isoforms in mouse livers at indicated ages. n = 3-4 mice/group. **(H)** Western blot analysis of ERAD proteins and known ERAD substrate IRE1α in P0 mouse livers. Two SEL1L antibodies (AB1, AB2) were used as shown **fig. S4F**. **(I)** Quantitation of western blot analysis of ERAD proteins and substrates in mouse livers. SEL1L from both antibodies were quantitated as average. Sex and age combined. n = 5-17 mice/group. Astericks, non-specific bands. Data are represented as means ± SEM. n.s., not significant. *p < 0.05; **p < 0.01; ***p < 0.001, ****p < 0.0001, Chi-square test for **(C)**; Two-way ANOVA followed by Tukey’s multiple-comparisons test for **(E and G)**; One-way ANOVA followed by Tukey’s post hoc test for **(I)**.

Interestingly, using the F1R1 PCR primer set spanning all 21 exons of SEL1L, we found that mouse *Sel1L* naturally expresses two mRNA isoforms in the liver (**Fig. 1, D to E**, Lanes 2 and 4). Sequencing revealed that these isoforms resulted from alternative splicing of exon 4: a predominant full-length isoform (*Sel1L-a*, ∼80%), and a shorter isoform (*Sel1L-b*) lacking 150 bp due to the use of an alternative “GU” splice donor site within exon 4 (arrow, **Fig. 1F**; illustrated in **Fig. 1D**). This splicing event removed amino acids 118–166, eliminating most of the FNII domain, including the C141Y mutation site (**Fig. 1F**). Notably, using the F2R2 primer set flanking exon 4 (**fig. S2A**), we found that despite large variations in overall *Sel1L* expression levels across mouse tissues (for example, a 5-fold difference between pancreas and spleen or lungs), the relative abundance of isoforms *a* and *b* remained consistent, with isoform *b* comprising ∼20% of total *Sel1L* transcript (**fig. S2, A to B**). Moreover, this splice pattern in the liver was unaffected by ER stress induced by tunicamycin injection (**fig. S2, C to D**).

Sequence analysis confirmed that no additional mutations were present in the *Sel1L* cDNA aside from the engineered C141Y mutation (data not shown). Remarkably, KI mice predominantly expressed the full-length *Sel1L-a* isoform, while the independently generated KI’ line primarily expressed shorter *Sel1L-b* isoform (**Fig. 1E**). In the KI’ line, the alternative splicing occurred at the same internal alternative “GU” splice donor site within exon 4, thereby excluding the pathogenic C141Y mutation (**Fig. 1F**). To further confirm this splicing pattern, we performed RT-PCR using the alternative primer pair flanking exon 4 (F2R2, **Fig. 1D**) and resolved the products via acrylamide gel electrophoresis to enhance separation of small fragments (**Fig. 1G**). While WT livers showed a 4:1 ratio of *Sel1L-a to -b*, KI and KI’ livers exclusively expressed *Sel1L-a* and *Sel1L-b*, respectively (**Fig. 1G**). Using an additional primer pair F3R3 spanning exons 1 and 6 (**fig. S3A**), we further confirmed the splicing patterns observed above (**fig. S3, B to E**). Notably, HET mice from both lines displayed intermediate isoform expression levels, falling between those of WT and the respective KI lines (**fig. S3, B to E**).

### Alternative splicing of *Sel1L* exon 4 rescues ERAD defects in KI’ mice

At the protein level, KI livers exhibited a near-complete loss of SEL1L expression, along with ∼ 60% reduction in the E3 ligase HRD1, ∼ 80% decrease in the ER lectin ERLEC1, and a ∼4-fold accumulation of the ERAD substrate IRE1α (*24*), consistent with ERAD dysfunction (**Fig. 1, H to I** and **fig. S4, A to C**). In contrast, KI’ livers restored SEL1L expression to ∼ 50% of WT levels despite harboring the same *Sel1L* C141Y mutation at the genomic level (**Fig. 1B**). These mice expressed a truncated SEL1L-B protein, ∼ 5 kDa smaller than the full-length isoform A, consistent with exon 4 skipping (**Fig. 1, H to I** and **fig. S4, D to E**). This was confirmed using two independent antibodies targeting distinct SEL1L epitopes (*18*) (**Fig. 1H** a**nd fig. S4F**). Expression of truncated SEL1L-B protein at ∼ 50% of WT levels was sufficient to restore HRD1 and ERLEC1 expression and to prevent IRE1α accumulation (**Fig. 1, H to I**), indicating preserved ERAD function in KI’ livers. HET mice from both lines showed intermediate SEL1L isoform expression between WT and their respective KI counterparts (**fig. S4, B to E**), consistent with a recessive inheritance model (*18*). Taken together, these findings demonstrate that alternative splicing of *Sel1L* exon 4 effectively rescues both ERAD defects and the lethality caused by the C141Y mutation *in vivo*.

### Disruption of the intronic splice donor enables alternative splicing in KI’ mice

To investigate the mechanism underlying the preferential use of an alternative splicing donor site within exon 4 in KI’ mice, we sequenced the genome region surrounding *Sel1L* exon 4. This analysis identified a single-nucleotide (T) deletion at the canonical “GT” splice donor site at the exon 4-intron 4 junction (**Fig. 2A**), likely introduced during the CRISPR/Cas9-mediated gene editing. To determine whether this disruption alone was sufficient to promote alternative splicing (**Fig. 2B**), we constructed a splicing minigene reporter (*25*) containing 168 bp exon 4 flanked by segments of 378 bp introns 3 and 463 bp intron 4, inserted between two GFP fragments (**Fig. 2C**).

**Fig. 2.**
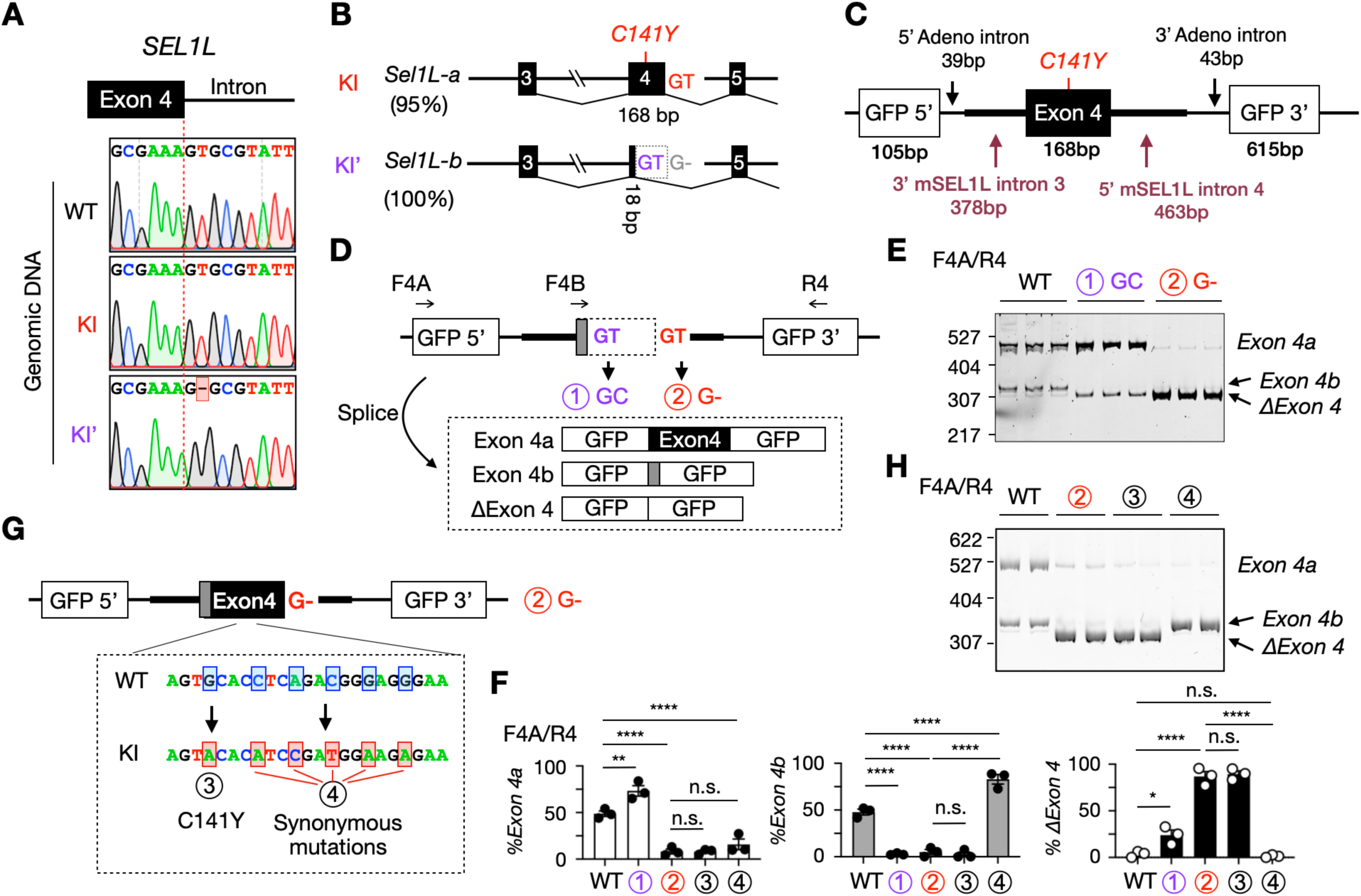
The mutation of major splicing donor enables the usage of an internal alternative splice donor. **(A)** Sanger sequencing results of SEL1L exon 4-intron 4 junction. Mutation was indicated by the red shaded box. **(B)** Schematic of the alternative splicing of Sel1L exon 4 with the percentage of isoforms in KI and KI’ mice indicated on the left. **(C)** Diagram of the splicing minigene reporter. **(D and G)** Diagrams of the indicated mutations in the minigene reporter construct, including primer design and the expected splicing products. **(E and H)** DNA PAGE analysis of the WT and mutated minigene reporters as indicated by using primer pair F4A/R4. **(F)** Quantitation of **(E)** and **(H)** as percent of exon 4 inclusion. n = 3 independent biological replicates. Data are represented as means ± SEM. n.s., not significant. *p < 0.05; **p < 0.01; ***p < 0.001, ****p < 0.0001, Two-way ANOVA followed by Tukey’s multiple-comparisons test for **(F)**.

RT-PCR using the F4A/R4 primer pair on GFP (**Fig. 2D**) revealed that the minigene containing WT *SEL1L* sequence yielded both full-length exon 4 and truncated exon 4b isoforms, consistent with utilization of both canonical and internal alternative splice donor sites (**Fig. 2E, quantitated in Fig. 2F;** sequencing data in **fig. S5A**). Mutation of the internal alternative splice donor site (mutation #1, GT → GC) abolished production of the exon 4b isoform, confirming its functionality (**Fig. 2E**, quantitated in **Fig. 2F**). Surprisingly, deletion of the canonical splice donor site (mutation #2, GT → G–) resulted in skipping of the entire exon 4 (*ΔExon 4*), rather than increase usage of the internal alternative site (**Fig. 2E**, quantitated in **Fig. 2F**). This result was corroborated using the F4B/R4 primer pair (**Fig. 2D**), which excludes the *ΔExon 4* product, and revealed minimal expression of the exon 4b isoform under these conditions (**fig. S5, B to C**).

This unexpected outcome prompted us to investigate whether additional sequence changes introduced during CRISPR/Cas9 editing – including the C141Y mutation and nearby synonymous mutations used for genotyping (**Fig. 2G**) – influence splice site selection. We introduced either the C141Y mutation (mutation #3, G → A) or the synonymous substitutions (mutation #4, C..A..C..G..G → A..C..T..A..A) into the minigene carrying mutation #2, which disrupts the canonical splice donor site (**Fig. 2G**). While the C141Y mutation did not alter the splicing pattern, the synonymous changes significantly enhanced usage of the alternative splice donor, resulting in increased expression of the exon 4b isoform (**Fig. 2H, quantitated in Fig. 2F**). These findings were further validated using F4B/R4 primer set (**fig. S5, B to E**). Together, these results suggest that loss of the canonical splice donor site creates a permissive context for exon skipping, while nearby exonic mutations – including synonymous changes – can promote use of an alternative splice donor site, likely by modulating exonic splicing regulatory elements.

### Pathophysiological significance of *Sel1L* alternative splicing

To assess the physiological relevance of *Sel1L* alternative splicing, we phenotypically characterized and compared KI and KI’ mice. While very few KI pups could survive past weaning (**Fig. 1C**), they were born – indicating that the mutation was not embryonic lethal – albeit at a reduced frequency of 16% at p0 (**Fig. 3A**). At birth, KI pups were indistinguishable from WT littermates in body weight (**Fig. 3B**), gross morphology (**Fig. 3C**), and tissue mass (**fig. S6, A to B**). However, by 12 hours postpartum, KI neonates lacked visible milk sacs (arrows, **Fig. 3C** and **fig. S6C**), indicative of impaired feeding. These phenotype parallels clinical reports of severe feeding difficulties in ENDI-A patients shortly after birth (*18*). Within the defined cohort, all KI pups died within 48 hours of birth (**Fig. 3D**), although rare survivors were observed in separate litters (5 out of 290 pups, **Fig. 1C**). Among these, 5 mice (4 males, 1 female) exhibited profound postnatal growth retardation (**Fig. 3E** and **fig. S6D**), mirroring the clinical presentation of ENDI-A (*18*). In contrast, KI’ mice were born at expected Mendelian ratios and showed normal growth comparable to WT and HET littermates (**Fig. 3E** and **fig. S6D**).

**Fig. 3.**
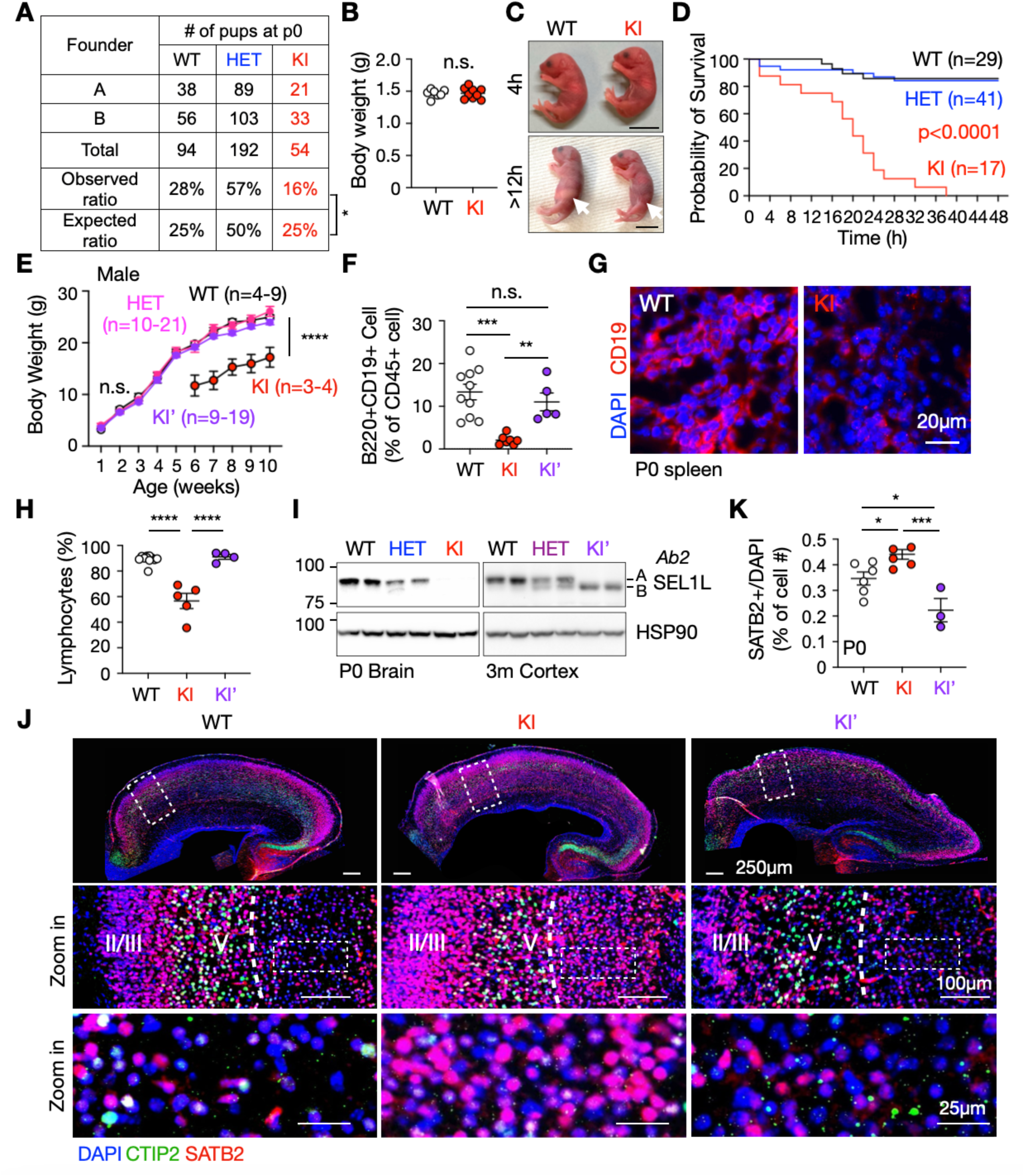
The alternative splicing of Sel1L exon 4 rescues developmental defects and B cell lymphopenia in SEL1L C141Y KI mice. **(A)** The survival rate of WT, HET and KI pups at postnatal day 0 (P0). **(B)** Body weight at P0. n = 7-9 mice/group. **(C)** The photo of pups at postnatal 4 hours (upper) and >12 hours (lower). **(D)** The survival curve of WT, HET and KI pups after birth. **(E)** Body weight growth of male WT, HET, KI’ and surviving KI mice for postnatal week 1 to week 10. **(F)** Quantification of B cells as a percentage of CD45⁺ peripheral blood mononuclear cells (PBMCs) at P0 by flow cytometry in **fig. S7, B to C**. n = 5-10 mice/group. **(G)** Immunofluorescence of CD19+ B cells in P0 mouse spleen with quantitation shown in **fig. S6F**. Red, CD19; Blue, DAPI. n = 3 mice/group. **(H)** Percentage of lymphocytes in the peripheral blood mononuclear cells (PBMCs) of WT, KI’ and surviving KI mice. Age and sex combined. n=4-11 mice/group. **(I)** Western blot analysis of SEL1L protein expression in mouse brains or cortex at indicated ages with quantitation in **fig. S8, B and D**. **(J and K)** Immunofluorescence **(J)** and quantitation **(K)** of SATB2+ Layer II/III cells number below the CTIP2+ Layer V in the cortex of p0 WT, KI and KI’ mice, the region as indicated by the Zoom in box. n = 3-6 mice/group. Green, CTIP2; Red, SATB2; Blue, DAPI. Data are represented as means ± SEM. n.s., not significant. *p < 0.05; **p < 0.01; ***p < 0.001, ****p < 0.0001, Chi-square test for **(A)**; Two-tailed t test for **(B)**; Mantel-Cox test for **(D)**; Two-way ANOVA followed by Tukey’s multiple-comparisons test for **(E)**; One-way ANOVA followed by Tukey’s post hoc test for **(F, H, K)**.

ENDI-A patients exhibit profound B cell lymphopenia (*18*), akin to mice with B cell-specific deletion of *Sel1L* (*26*). Similarly, KI neonates exhibited marked reductions in mature B cells in peripheral blood and spleen at p0, as measured by flow cytometry (**Fig. 3F** and **fig. S7, A to B**), histology (arrows, **fig. S6E**), and immunofluorescence staining using mature B cell markers CD19 and B220 (**Fig. 3G**, and **fig. S6F**). Surviving adult KI mice also showed significantly reduced lymphocytes in peripheral blood mononuclear cells (PBMCs) (**Fig. 3H**) and a notable expansion of circulating neutrophils (**fig. S6G**), a phenotype commonly observed in B cell-deficient models (*27*). By contrast, KI’ mice showed normal circulating B cell numbers at birth (**Fig. 3F** and **fig. S7C**) and puberty (**fig. S7, D to E)**, with no elevation in neutrophils (**Fig. S6G**).

Given that ENDI patients also present with intellectual disability (*17, 18*), we next examined cortical development at p0. As observed in the liver (**Fig. 1H**), KI brains showed near-complete loss of SEL1L expression, reduced HRD1 protein levels (by ∼ 70%), and ERAD dysfunction – all of which were restored in KI’ mice (**Fig. 3I, fig. S8, A to D**). Cortical development occurs during embryogenesis and results in the formation of six distinct layers arranged in an inside-out pattern from Layer VI to Layer I (*28*). In WT brains, immunofluorescence staining revealed well-organized cortical layering, with clearly defined Layer ΙΙ/ΙΙΙ (SATB2-positive) and Layer V (CTIP2-positive) neurons (**Fig. 3, J to K**). In contrast, KI mice exhibited disrupted cortical architecture, including ectopic localization of SATB2-positive cells below Layer V without obvious cell loss, indicative of impaired neuronal migration or cortical specification (**Fig. 3, J to K**). Interestingly, KI’ mice displayed a normalized SATB2+ cell distribution and restored laminar organization (**Fig. 3, J to K**). Moreover, we observed minimal UPR activation as indicated by the absence of ER chaperone induction (BiP and PDI, **fig. S9A**) and lack of UPR sensor activation, including IRE1α and PERK phosphorylation and *Xbp1* mRNA splicing, in the brains of KI pups (**fig. S9, B to D**). Collectively, these findings establish pathogenic nature of the *SEL1L* C141Y variant and demonstrate that exon 4 skipping via alternative splicing functionally rescue the lethal and developmental defects associated with the mutation *in vivo*.

### ASO-mediated exon skipping rescues ERAD defects in patient-derived fibroblasts

Prompted by serendipitous observations in mice, we next explored whether an exon-skipping strategy could similarly be applied to human cells harboring the C141Y mutation. Interestingly, sequence alignment revealed that the internal “GU” splice donor site found in murine exon 4 is not conserved in humans or other species (**Fig. 4A**), consistent with the exclusive expression of the full-length *SEL1L-A* isoform in human tissues (**Fig. 1E**). To test the feasibility of exon skipping in this context, we treated fibroblasts from a SEL1L C141Y patient with a panel of 25-nucleotide-long ASOs targeting exon 4 and adjacent intronic sequenes. RT-PCR using F5R5 flanking primer pair (**Fig. 4C**), followed by Sanger sequencing, revealed the emergence of two alternative splice variants in addition to the full-length isoform A: isoform *B,* which lacked the entire exon 4 and was induced by ASO1 and ASO3-5; and isoform *C* detected with ASO6-7, which lacked the first 63 bp of exon 4, likely due to the disruption of the intron 3 splicing acceptor and activation of a cryptic splice site within exon 4 (**Fig. 4, C to D** and **fig. S10**). Among all the tested ASOs, ASO1 – targeting the exon 4-intron 4 junction – was the most effective, inducing exon 4 skipping in approximately 60% of transcripts in both WT and patient fibroblasts (**Fig. 4, C to E** and **fig. S11A**). Similar dose-dependent exon-skipping results were observed in WT HEK293T cells (**fig. S11B**).

**Fig. 4.**
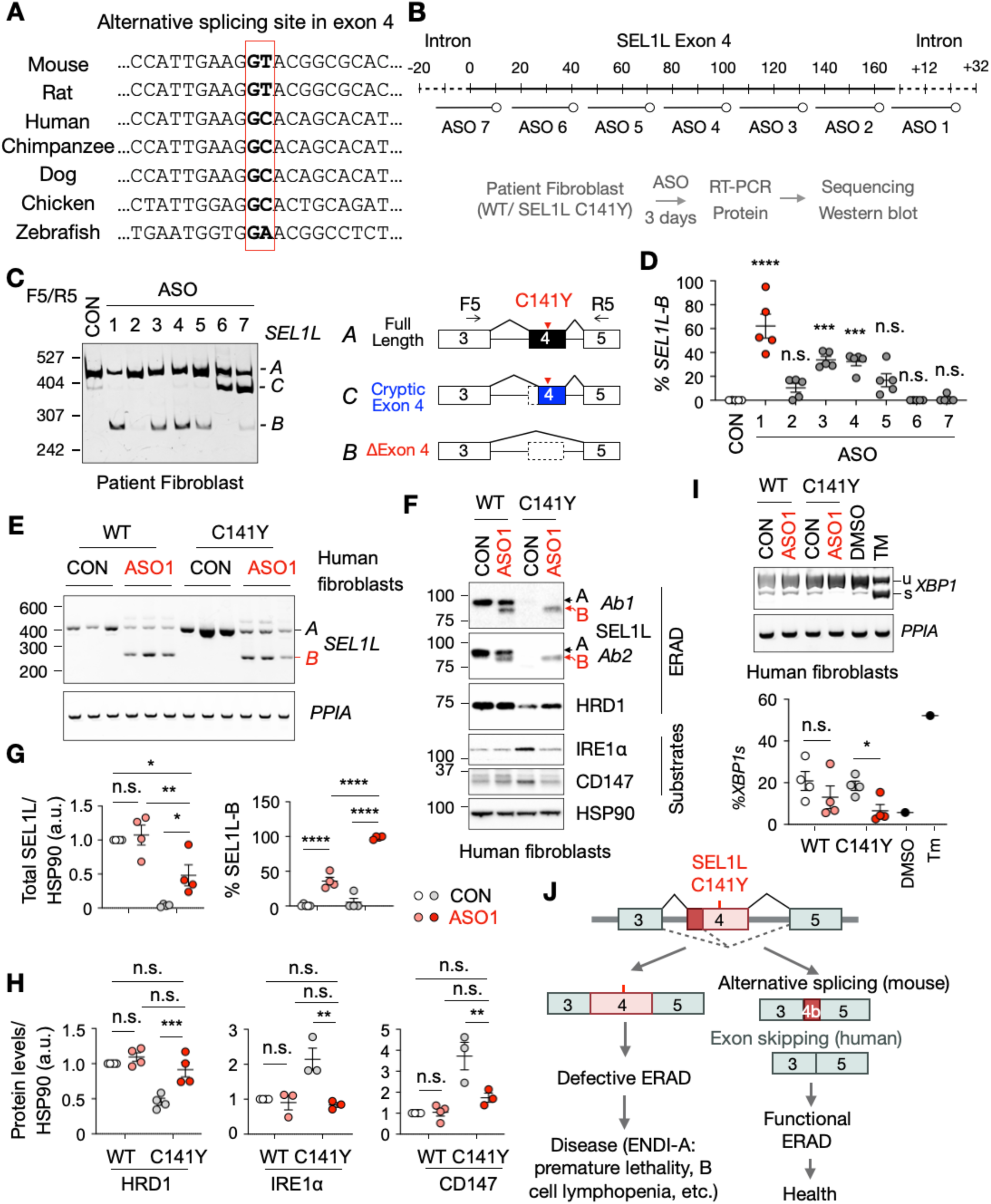
ASO-mediated exon skipping rescues ERAD defects in C141Y patient-derived fibroblasts. **(A)** Sequence alignment of the alternative splice site within *Sel1L* exon 4 across different species. **(B)** The experimental design of the ASOs screen targeting SEL1L Exon 4. **(C and D)** DNA PAGE analysis **(C)** and quantitation **(D)** of ASOs treatment at 10 μM for 3 days in C141Y patient fibroblasts, with a diagram showing corresponding products. C, Control ASO. Red arrowhead indicates the location of SEL1L C141Y mutation. n = 5 independent biological replicates. Statistics indicates the comparison between control ASO and ASO 1-7 targeting SEL1L exon 4 and introns 3 and 4. **(E)** DNA agarose gel electrophoresis analysis of ASO1 treatment at 10μM for 3 days in human WT and C141Y patient fibroblasts. Quantitation shown in the **fig. S11A**. n = 5-6 independent biological replicates. **(F to H)** Western blot analysis **(F)** and quantitation **(G and H)** of ASO1 treatment at 10μM for 3 days in human WT and C141Y patient fibroblasts. Two SEL1L antibodies (AB1, AB2) were used as shown **fig. S4F**. n = 3-4 independent biological replicates. **(I)** Electrophoresis analysis of XBP1 splicing of ASO1 treatment at 10μM for 3 days in human WT and C141Y patient fibroblasts, with quantitation below. DNA PAGE gel for the *XBP1* splicing and agarose gel for the internal control *PPIA*. n = 3 independent replicates for each group. n = 1 for positive controls. u, unspliced. s, spliced. **(J)** Schematic model illustrating rescue of the SEL1L C141Y mutation through alternative splicing modulation in mice and humans. Data are represented as means ± SEM. n.s., not significant. *p < 0.05; **p < 0.01; ***p < 0.001, ****p < 0.0001, One-way ANOVA followed by Tukey’s post hoc test for **(D)**; Two-way ANOVA followed by Tukey’s multiple-comparisons test for **(G and H),** Multiple t-tests for **(I)**.

ASO1 treatment significatntly inceased expression of the truncated SEL1L-B protein in WT fibroblasts, accounting for ∼ 40% of total SEL1L protein, and rescued SEL1L expression in C141Y patient fibroblasts (**Fig. 4F** and quantitated in **Fig. 4G)**. Neither SEL1L C141Y mutation nor ASO treatment altered *SEL1L* mRNA levels (**fig. S11C**), suggesting a post-transcriptional effect. Consistent with the role of SEL1L in stabilizing HRD1 (*29*), ASO1 treatment restored HRD1 protein levels and rescued ERAD function, as evidenced by decreased accumulation of known ERAD substrates including IRE1α (*24*) and CD147 (*30*) (**Fig. 4F** and quantitated in **Fig. 4H**), and reduced *XBP1* mRNA splicing (**Fig. 4I**).

To further validate these findings, we employed CRISPR/Cas9 to engineered KI HEK293T cells. While introduction of the C141Y or C127Y mutation abolished SEL1L protein expression, complete deletion of the FNII domain produced a truncated SEL1L protein (∼5 kDa smaller than WT) that preserved normal ERAD function, as reflected by unchanged IRE1α levels (*18*) (**fig. S11, D to E**). These results indicate that although disulfide bonds within the FNII domain are cirtical for SEL1L function, the domain itself is dispensable for ERAD function. Taken together, these findings demonstrate that ASO-mediated exon skipping restores functional SEL1L and rescues ERAD defects in patient-derived fibroblasts, highlighting its therapeutic potential for patients with SEL1L defects.

## DISCUSSIONS

The SEL1L-HRD1 ERAD pathway is a highly conserved protein quality control mechanism that plays a central role in maintaining ER homeostasis (*14*). We recently identified biallelic mutations in *SEL1L* and *HRD1* in patients with ENDI and ENDI-A syndromes (*17, 18*), implicating ERAD dysfunction in the pathogenesis of these rare but severe genetic disorders. Using a KI mouse model, we established the pathogenicity of the SEL1L C141Y variant located in the fibronectin type II (FNII) domain. Moreover, although this residue is essential for SEL1L protein stability, we found that the FNII domain itself is dispensable for SEL1L function and ERAD activity. This unexpected finding, rooted in the phenotypic discrepancies among three independently generated KI mouse lines carrying the same mutation, led to the discovery of naturally occurring alternative splicing as a mechanism of functional rescue (**Fig. 4J**). Thus, our study not only defines the molecular pathology of the C141Y mutation but also highlights splicing modulation as a promising therapeutic strategy for correcting ERAD defects.

The SEL1L FNII domain is conserved across vertebrates and is also found in proteins such as fibronectin and tissue-type plasminogen activator/coagulation factor XII (*18, 22*), but it is absent in yeast SEL1L counterpart Hrd3. This pattern suggests that the domain may have been evolutionarily acquired during vertebrate evolution through exon shuffling (*31*). In rodent SEL1L, this domain is encoded by exon 4, which undergoes alternative splicing to generate isoform diversity – a feature not observed in humans. This rodent-specific splicing event may provide an added layer of control over SEL1L function. Although the precise role of the FNII domain remains incompletely define, our data indicate that SEL1L-B, which lacks this domain, is less stable than the full-length protein, suggesting a role in stabilizing SEL1L. In addition, a recent study implicated SEL1L in collagen turnover and proposed that its FNII domain may contribute to collagen binding (*32*). While we speculate that the FNII domain of SEL1L may be involved in substrate recruitment, further studies will be needed to delineate its precise function and broader biological relevance.

Although overexpression of SEL1L or HRD1 has not been directly linked to toxicity *in vivo*, excessive ERAD activity has been associated with cancer progression (*33–35*), dampening enthusiasm for long-term gene replacement approaches. Similar challenges exist in other dosage-sensitive disorders; for example, loss-of-function mutations in *MECP2* cause Rett syndrome, whereas duplications of the same gene result in a separate neurodevelopmental disorder (*36, 37*). In this context, our study shows that exon 4 skipping — whether naturally occurring in mice or induced by ASOs in human cells — can restore ERAD function (**Fig. 4J**). By generating a truncated yet functional SEL1L protein that bypasses the fatal mutation, ASO-mediated exon skipping overcomes the instability of mutant SEL1L and avoids potential toxicity associated with gene overexpression. While our data showed that this approach could rescue ERAD function in patient-derived fibroblasts *in vitro*, whether this approach can prevent perinatal lethality and developmental defects *in vivo* requires further careful investigations.

The clinical success of ASO therapies, such as eteplirsen for Duchenne muscular dystrophy and approved treatments for spinal muscular atrophy and amyotrophic lateral sclerosis, underscores their utility in correcting genetic defects by restoring partially functional proteins with high specificity and minimal off-target effects (*21, 38, 39*). Intrathecal delivery of ASOs has shown promise in treating rare neurodevelopmental disorders, with substantial improvements in seizures and cognitive outcomes when administered early (*40–43*). To date, ASOs have not been applied to diseases involving ERAD dysfunction. Our findings establish a proof-of-concept for this approach, demonstrating that targeted exon skipping can restore proteostasis in the setting of a pathogenic *SEL1L* mutation. Given the essential role of SEL1L-HRD1 ERAD in cellular homeostasis (*3, 4, 14, 16*), this strategy may hold promise not only for *SEL1L*-related disorders but also for a broader range of protein misfolding diseases.

## MATERIALS AND METHODS

### Study Design

This study investigates the pathogenicity of the SEL1L C141Y mutation and evaluates the therapeutic potential of alternative splicing *in vivo* and via antisense oligonucleotide (ASO)-mediated exon skipping in patient-derived fibroblasts. All animal procedures were approved by the Institutional Animal Care and Use Committee of the University of Michigan Medical School (PRO0008989 and PRO00010658), and University of Virginia Medical School (Protocol No. 4459) and in accordance with the National Institutes of Health (NIH) guidelines. Sample sizes were determined on the basis of prior literature and power calculations to ensure statistically significant results while minimizing the number of animals used in accordance with ethical guidelines. Blinding was applied during running flow cytometry, complete blood count, and histological analysis. Experiments were replicated at least 3 times to ensure reproducibility, and sample sizes for each study are reported in the figure legends. Data inclusion criteria required samples to meet experimental conditions without visible contamination or procedural errors, and exclusion criteria involved procedural inconsistencies, or abnormal physiological conditions in animals (e.g. severe dystocia in breeder female for pup survival experiment). Outliers were assessed both visually using Q-Q plots and statistically using the Shapiro-Wilk or Kolmogorov-Smirnov tests (for sample sizes greater than three), with a significance threshold of α = 0.05. Outliers linked to experimental or methodological errors were excluded, while those reflecting biological variation were retained. For expanded Materials and Methods used in this article, please see the Supplementary Materials.

### Mice

The *SEL1L^C141Y^* knock-in (KI) mice (human SEL1L p.Cys141 is equivalent to mouse SEL1L p.Cys137) were generated at the University of Michigan Molecular Genetics Core using the CRISPR-Cas9 technology. The two single guide RNAs (sgRNAs) were designed by using computer algorithm (http://crispor.tefor.net), targeting mouse genome *SEL1L* exon 4 where the mutation is located: sgRNA1: 5’-ATGAGTGCACCTCAGACGGG-3’; sgRNA2: 5’- AAGTGCGTATTGTTCAAGTG -3’. The sgRNAs were synthesized using the Synthego sgRNA synthesis Kit per manufacturer’s protocol, tested and confirmed in fertilized eggs. A donor DNA carrying mouse *Sel1L* cDNA 410G>A mutation and additional silent mutations was designed to mediate homology-directed repair (HDR): 5’- CACGGGGAGCCCTGCCACTTCCCTTTCCTTTTCCTGGATAAGGAGTATGATGAGTACACATCCGAT GGAAGAGAAGATGGCAGACTGTGGTGTGCCAC AACCTATGACTACAAGACAGATGAGAAGTGGGGCTTCTGCGAAAGTGCGTATTGTTCAAG TGCACGCCCTGTGCTTTAGGGCAGCATTTGGAAGGCATTTTC-3’. The donor DNA was synthesized as Ultramer dsDNA by Integrated DNA Technologies, Inc. A mixture of Cas9 protein (Sigma), sgRNAs, and donor DNA was microinjected into fertilized mouse eggs on the C57BL6/J background. The injected zygotes were then transferred into pseudopregnant females. The existence of desired mutations was further confirmed by Sanger sequencing in three independent founders. The founders were then bred separately to WT C57BL6/J mice to obtain F1 heterozygous *SEL1L^C141Y^*heterozygous mice. F1 heterozygous *SEL1L^C141Y^* mice were inter-crossed to generate homozygous *SEL1L^C141Y^* KI mice and its WT and heterozygous littermates. For survival curve, pregnant female and the litter size were checked every 4 hours. Pup tails were collected for genotyping. Age- and gender-matched littermates were maintained in a temperature-controlled room on a 12-h light/dark cycle and used in all studies. For *in vivo* ER stress induction, 12 weeks old male WT mice were used for UPR activation by DMSO or tunicamycin treatment at 1mg/kg for 4 hours via intraperitoneal injection.

### Statistical analysis

Statistics tests were performed in GraphPad Prism version 10.0 (GraphPad Software). Unless indicated otherwise, values are presented as mean ± standard error of the mean (SEM). All experiments have been repeated at least three times and/or performed with multiple independent biological samples from which representative data are shown. Statistical differences between the groups were compared using the unpaired two-tailed Student’s *t*-test for two groups, one-way ANOVA followed by Tukey’s post hoc test, two-way ANOVA followed by Tukey’s multiple-comparisons test for multiple groups, Chi-square test for contingency table, Mantel-Cox test for survival curve. *P* < 0.05 was considered statistically significant.

## AUTHOR CONTRIBUTIONS

H.H.W. designed and performed most experiments; Z.W., H.W., Z.J.L., and L.L.L helped with the mouse survival curve and *in vivo* experiments; L.J. designed and W.G. helped with the neurological experiments; S.K.M. and M.N.K. provided guidance and helped with the exon skipping and alternative splicing experiments; S.S. and L.Q. directed the study and designed experiments. H.H.W. and L.Q. wrote the manuscript; all authors commented on and approved the manuscript.

## ACKNOWLEDGEMENTS

We acknowledge Drs. Fowzan S Alkuraya, Denisa Weis, Nicola Brunetti-Pierri, and members in the Qi, Sun, Arvan and Kuyumcu-Martinez laboratories for reagents, technical assistance, insightful discussions, and constructive comments on the manuscript. We acknowledge Ellie Sherman in the Jiang lab for the technical assistance on the brain-related experiments. We acknowledge University of Michigan Molecular Genetics Core for the generation of SEL1L C141Y KI mice, the University of Virginia School of Medicine Research Histology Core, Flow Cytometry Core, and Biorepository and Tissue Research Facility Core for their help. H.H.W. is supported by Alzheimer’s Association Research Fellowship 25AARF-1375486. This work was supported by Additional Ventures Single Ventricle Research Fund 1048010, and in part by 5R01HL157780-03 and 1R01HL175488-01 (M.N.K-M); R01DK128077, R01DK132068 (S.S.); Alzheimer’s Association 24AARGD-NTF-1187603, R35GM130292, 1R01AG089640 (L.Q.), and 1R01NS138119 (L.Q. and S.S.).

## Data and materials availability

The materials and reagents used are either commercially available or available upon the request.

