## Supplementary Materials for "Functional rescue of a fatal ERAD mutation via alternative splicing"

**List of Supplementary Materials**

Materials and Methods

Fig S1 to S11

### MATERIALS AND METHODS

#### Human Liver Tissue

Deidentified frozen normal human liver tissue was acquired through Biorepository and Tissue Research Facility Core at the University of Virginia.

#### Genotyping, PCR and sequencing

Mice were routinely genotyped using PCR of genomic DNA samples obtained from tails or ears with the following primer pairs:

*Sel1L*$^{C141Y}$ allele for *SEL1L*$^{C141Y}$ KI mice (**fig. S1, B to C**):

F: 5'- AGTACACATCCGATGGAAGAGAAG-3';

R: 5'-GAAAATGCCTTCCAAATGCTGC-3';

*Sel1L* wildtype allele (**fig. 1S, B to C**):

F: 5'- AGTGCACCTCAGACGGGAGGG-3';

R: 5'- GAAAATGCCTTCCAAATGCTGC-3';

Mouse tail or ear genomic DNA sample or cell genomic DNA was used for sanger sequencing analysis by using following primers for PCR and sequencing:

Sequencing for *mSel1L* genomic DNA Exon 4 and intron 4 (**Fig. 2A**):

F: 5'- CTTAAGAACTCAAAGTCTACACTAAGTCT-3';

R: 5'- CAGCTTGCCTCAAGGGTTTACAGAA-3';

#### Histology and immunofluorescence

Adult mice were anesthetized and perfused with 20 ml of PBS followed by 40 ml of 4% paraform-aldehyde in 0.1 M PBS pH 7.4 for fixation. Tissues were dissected out and fixed overnight in 4% paraformaldehyde in PBS at 4ºC. For hematoxylin and eosin (H&E) staining, samples were de-hydrated, embedded in paraffin, and stained at the Research Histology Core at the University of

Virginia School of Medicine. Images were captured using Aperio ScanScope CS System and Aperio ImageScope software.

For immunofluorescence staining, postnatal P0 pups were anesthetized with hypothermia. Then, pup spleen and brain were dissected followed by overnight fixation in 5mL 4% paraformaldehyde in 0.1 M PBS pH 7.4 under 4ºC. The tissues were then transferred to 30% sucrose in PBS solution for overnight incubation under 4ºC and then embed in OCT for cryosection. The tissues were sectioned at 8-15μM. The samples were blocked with 5% normal donkey serum, 0.3% Tween-20 PBS solution for 1 hour and then incubated with primary antibody diluted in 5% normal donkey serum, 0.3% Tween-20 PBS solution for overnight at 4°C: anti-CD19 (Cell Signaling Technology, #90176S, 1:100); anti-SATB2 (Abcam, ab51502, 1:20); anti-CTIP2 (Abcam, ab18465, 1:100). The samples were washed with PBS for 5 minutes x 3 times at room temperature, and then incubated with secondary antibody for 1 hour at room temperature: Alexa Fluor 555-conjugated donkey anti-rabbit IgG (Jackson ImmunoResearch, 711-565-152, 1:500), Alexa Fluor 488-conjugated goat anti-rat IgG (Invitrogen, A-11006, 1:500), Alexa Fluor 555-conjugated donkey anti-mouse IgG (Invitrogen, A32773, 1:500). After 5 minutes x 3 times washing with PBS at room temperature, the slides were mounted in ProLong™ Diamond Antifade Mountant with DAPI (Fisher Scientific P36971). The samples were imaged by Leica DMi8 THUNDER Imager.

#### Plasmids

The following plasmids were used in the study: pcDNA5-GFP-IL7R-XbaI mutated minigene construct was a kind gift from Dr. Muge N Kuyumcu-Martinez Lab (*1*). Then, pcDNA5-GFP-SEL1Lexon4 were generated by GenScript Biotech, by replacing IL7 exon in the pcDNA5-GFP-IL7-XbaI mutated minigene construct with mouse SEL1L exon 4 and the proximal introns (348 bp upstream and 462 bp downstream). Alternative splicing donor site mutation and exon 4 splicing donor site mutation were generated using the above mentioned pcDNA5-GFP-SEL1Lexon4 minigene construct. All plasmids were validated by DNA sequencing. The primers used for mutagenesis in **Fig. 2D** and **2G** are:

Mutagenesis primers for alternative splicing donor site (Mutation 1):

F 5'-CCATTGAAGGCACGGCGCACGGGGA-3'

R 5'-GCGCCGTGCCTTCAATGGCAGTCAAGA-3'

Mutagenesis primers for Exon 4 canonical splicing donor site mutation (Mutation 2):

F 5'-CTTCTGCGAAAGGCGTATTGTTCAAGTGGGG-3'

R 5'-AACAATACGCCTTTCGCAGAAGCCCCACTTCT-3'

Mutagenesis primers for Exon 4 C137Y mutation (Mutation 3):

F 5'- GTATGATGAGTACACCTCAGACG-3'

R 5'- CGTCTGAGGTGTACTCATCATAC-3'

Mutagenesis primers for Exon 4 synonymous mutations (Mutation 4):

F 5'- GATGAGTGCACATCCGATGGAAGAGAAGATGGCAGACTGTGG-3'

R 5'- GCCATCTTCTCTTCCATCGGATGTGCACTCATCATACTCC-3'

### RNA preparation and RT-PCR

Total RNA was extracted from cell or tissues using TRI Reagent and BCP phase separation reagent (Molecular Research Center, TR 118). For RT-PCR analysis the following primer sequences were used:

*mSEL1L* Full length (F1/R1 used for **Fig. 1E**) F: 5'- ATGCAGGTCCGCGTCAGGCTGTCGTT-GCTGCT-3'; R: 5'- CTACTGTGGTGGCTGCTGCTCTGG-3';

*hSEL1L* Full length (F1'/R1' used for **Fig. 1E**) F: 5'- GCGGCTAGCATGCGGGTCCGGA-TAGGGCT-3'; R: 5'- GCGAAGCTTTTACTGTGGTGGCTGCTGCTCTG-3';

*mSEL1L* Exon4 for acrylamide gel (F2/R2 used for **Fig. 1G and fig. S2, A and C**) F: 5'- AGCAA-GACCTACGAAGAACT-3'; R: 5'- GAAGGTACCGATATGCTTCTCTCT-3';

*mSEL1L* Exon4 for agarose gel (F3/R3 used for **fig. S3**): F: 5'-ATGCAGGTCCGCGTCAGGCTGTCGTTGCTGCT -3';

R: 5'- CTACTGTGGTGGCTGCTGCTCTGG -3';

Minigene Exon4-1 (F4A/R4 used for **Fig. 2, E and H**): F: 5'- TGGTGAGCAAGGGCGAGG-3'; R: 5'- CGTCCTTGAAGAAGATGGTGCG-3';

Minigene Exon4-2 (F4B/R4 used for **fig. S5, B and F**): F: 5'- GCGAGTCTTGACTGCCATTGAAG-3'; R: 5'- CGTCCTTGAAGAAGATGGTGCG-3';

*hSEL1L* Exon4 (F5/R5 used for **Fig. 4C, 4E and fig. S11B**): 5'- GGGGAAAGTGTCACAGAAGA-TATCAG-3'; R: 5'- GACACTCTCTCCAGGGCTTTG-3';

*mXbp1s*: F: 5'- ACGAGGTTCCAGAGGTGGAG-3'; R: 5'- AAGAGGCAACAGTGTCAGAG-3';

*hXBP1s*: F: 5'- GAATGAAGTGAGGCCAGTGG-3'; R: 5'- ACTGGGTCCTTCTGGGTAGA-3';

*mL32* F: 5'- GAGCAACAAGAAAACCAAGCA-3'; R: 5'- TGCACACAAGCCATCTACTCA-3';

*hPPIA* F: 5'- GGCAAATGCTGGACCCAACACA-3'; R: 5'- TGCTGGTCTTGCCATTCCTGGA-3';

RT-PCR products were analyzed by 0.8%-1.5% agarose electrophoresis or 5% polyacrylamide gel electrophoresis gel using CBS Scientific Adjustable Height Vertical Gel System.

### Western blot and antibodies

Mouse tissues or cells were harvested and snap-frozen in liquid nitrogen. The proteins were extracted by sonication in NP-40 lysis buffer (50 mM Tris-HCl at pH7.5, 150 mM NaCl, 1% NP-40, 1 mM EDTA) with protease inhibitor (Sigma), DTT (Sigma, 1 mM) and phosphatase inhibitor cocktail (Sigma). Lysates were incubated on ice for 30 min and centrifuged at 16,000 g for 10 min. Supernatants were collected and analyzed for protein concentration using the Bio-Rad Protein Assay Dye (Bio-Rad). 20-50 μg of protein were denatured at 95°C for 5 min in 5x SDS sample buffer (250 mM Tris-HCl pH 6.8, 10% sodium dodecyl sulfate, 0.05% bromophenol blue, 50% glycerol, and 1.44 M β-mercaptoethanol). Protein was separated on SDS-PAGE, followed by electrophoretic transfer to PVDF (Fisher Scientific) membrane. The blots were incubated in 2%

BSA/Tri-buffered saline tween-20 (TBST) overnight at 4°C with primary antibodies specifically for: HSP90 (Santa Cruz, #sc-13119, 1:5,000), SEL1L (home-made, 1:10,000) (*2*), SEL1L (Abcam, ab78298, 1:1,000), HRD1 (Proteintech, #13473-1, 1:2,000), CD147 (Proteintech, #11989-1, 1:3,000), IRE1α (Cell Signaling, #3294, 1:2,000), ERLEC1 (Abcam, #ab181166, 1:5,000), BiP/GRP94 (Abcam, #ab21685, 1:5,000), PDI (Enzo, #ADI-SPA-890, 1:5,000), PERK (Cell Signaling, #3192, 1:5000). Membranes were washed with TBST and incubated with secondary antibodies, either HRP conjugated (Bio-Rad, 1:10,000), anti-Rabbit IgG TrueBlot HRP (Rockland, #18-8816-33, 1:500) or anti-Mouse IgG TrueBlot-HRP (Rockland, #18-8817-31, 1:500) at room temperature for 1h for ECL chemiluminescence detection system (Bio-Rad) development. Phos-tag-based western blot analysis was performed as previously described (*3*). Band intensity was determined using ImageJ FIJI software.

#### Antisense Oligonucleotide (ASO) Design and Treatment

Morpholino ASO were either designed by Morpholino Gene Tool, LLC or by the authors, and synthesized by Morpholino Gene Tool, LLC. Standard Control Oligo (CCTCTTACCTCAG-TTACAATTTATA) were purchased from Morpholino Gene Tool, LLC. Morpholino ASO were dissolved in the original vial with sterilized $H_2O$ at 1mM. The human fibroblasts or HEK293T cells were cultured at 80-100% confluency and treated with 10μM Morpholino oligo and 6 μM/mL Endo-Porter (Morpholino Gene Tool, LLC, OT-EP-PEG-1) in 10% serum DMEM medium. The cells were collected after 1 or 3 days for RNA or protein analysis, respectively. The ASO sequences are:

ASO1: TGCCTCCTACTGAGCAATACTTACT

ASO2: AAAAGCCCCACTTTTCATCTGCTTT

ASO3: CATAGGTTGTAGCACACCACAGTCT

ASO4: CTTCCCTCCCATCTGATGTATATTC

ASO5: ACTCCTTATCTAGGAAAAGAAAAGG

ASO6: GGCAGGGCTCCCCATGTGCTGTGCC

ASO7: GGCGGTCAAAGCTGGAATGACAAGA

### Complete Blood Count (CBC), Flow Cytometry, Intracellular Staining, and Antibodies

Peripheral blood was collected from surviving KI mice via facial vein into EDTA-coated tubes and complete blood count was analyzed by University of Virginia Center for Comparative Medicine (CCM) Core. Flow cytometric analysis of peripheral blood mononuclear cells (PBMCs) was performed as we described previously (*4, 5*). In brief, PBMCs were isolated from blood treated with anti-coagulation reagent and red blood cell lysis buffer for 5 minutes at room temperature. The PBMCs were then stained with ZombieNIR Fixable Viability dye (BioLegend 423106) diluted as 1:400 in PBS-$Ca^{2+}$ solution for 10 minutes at room temperature. CD16/CD32 Rat anti-Mouse antibody was then added as 1:100 to PBMCs for blocking at 4ºC for 5 minutes. The PBMCs were then incubated with 1:100 fluorochrome-conjugated antibody against anti-CD45 (30-F11), anti-CD45R/B220 (RA3-6B2), Anti-CD19 (6D5) (BioLegend or eBioscience) at 4ºC for 30 minutes, followed by 2 washes with Flow cytometry (FACS) buffer (5% serum PBS-Ca solution) under 4ºC. The samples were then fixed with 2% paraformaldehyde under room temperature for 20 minutes, followed by 2 washes. The samples were stored at 4ºC overnight and analyzed on Cytek Aurora Northern Lights Spectral Flow Cytometer at the University of Virginia Flow Cytometry Core. The .fcs file were analyzed on FCS Express 7 Research.

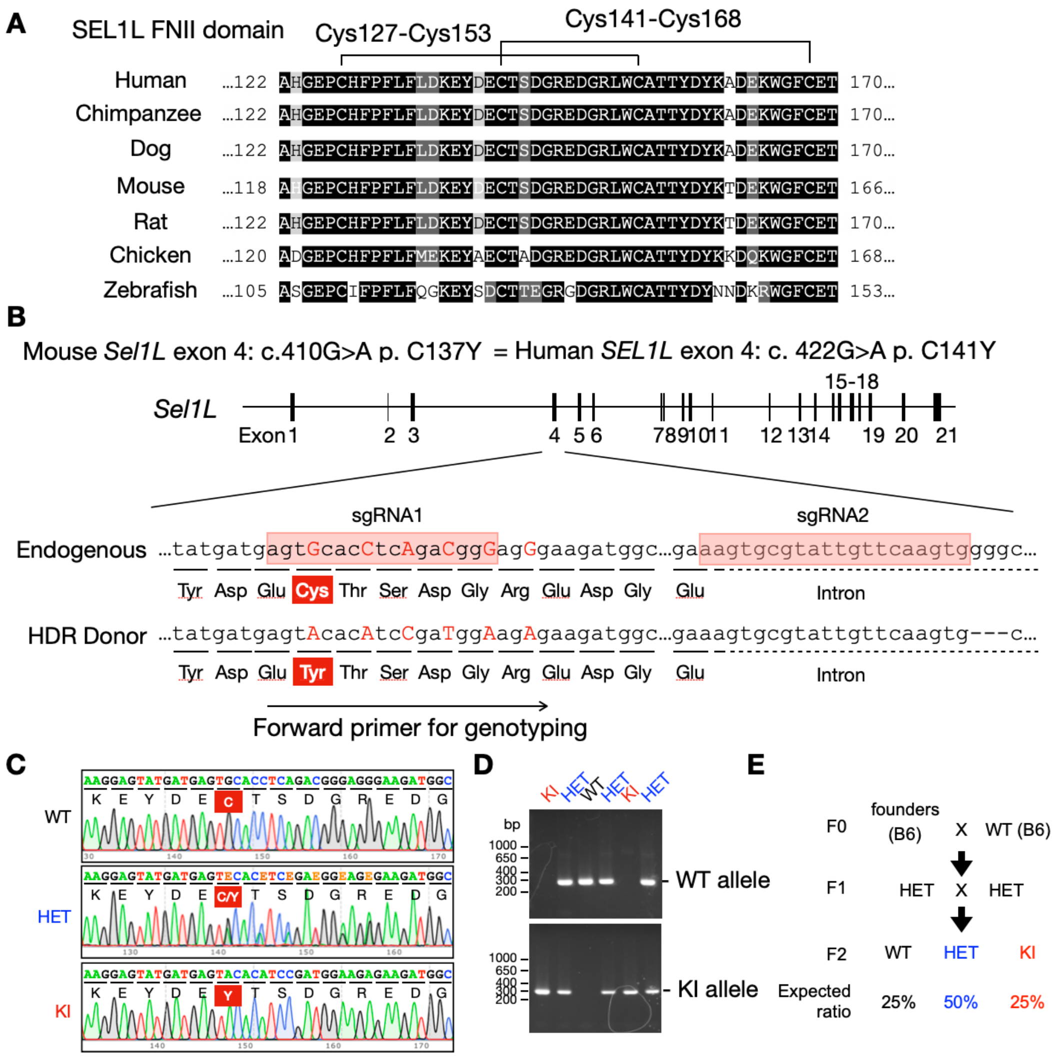


**Fig. S1. The generation of SEL1L C141Y mutation knock-in (KI) mouse**. **(A)** Sequence alignment of the FNII domain across vertebrate species. Lines indicate conserved disulfide bond pairs. **(B)** Diagram of sgRNAs and homology-directed repair (HDR) donor used to generate SEL1L C141Y knock-in mice via CRISPR-Cas9 technology. Silent mutations were introduced to facilitate genotyping, and the "GGG" sequence was deleted in the HDR donor (indicated as "– – –") to disrupt the PAM site. Forward primer for genotyping is indicated below the HDR donor sequence. **(C)** Sanger sequencing confirming the introduction of the SEL1L C141Y mutation along with surrounding silent mutations. **(D)** DNA agarose gel electrophoresis for genotyping of SEL1L C141Y knock-in mice. **(E)** Breeding strategy for SEL1L C141Y KI mice. Each established founder line was bred independently.

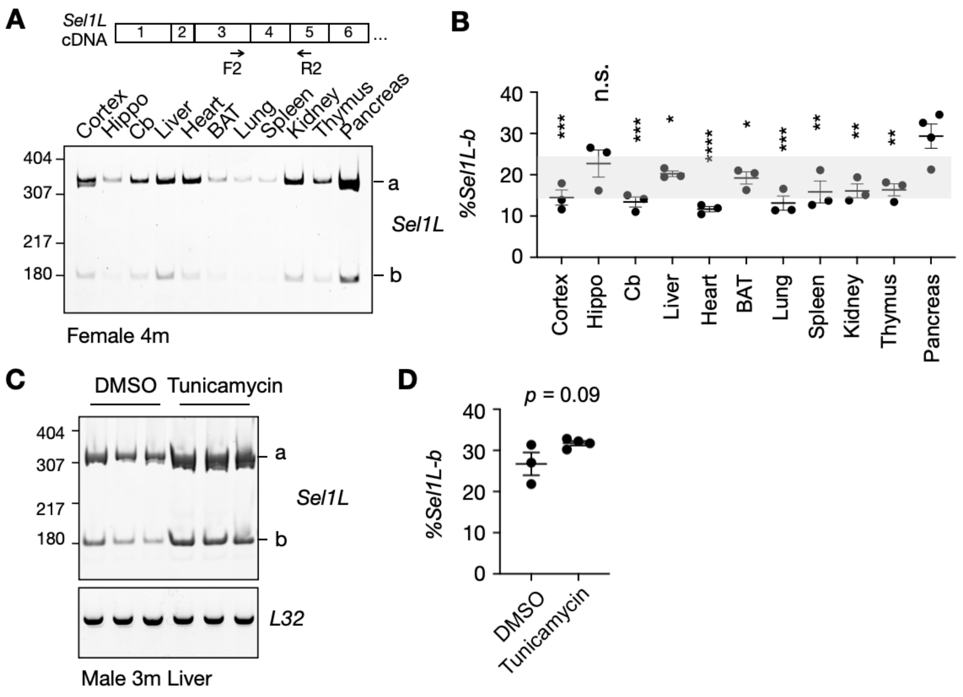


**Fig. S2. The alternative splicing of Sel1L occurs across different tissues in mice and is not affected by ER stress. (A to B)** Diagram and DNA polyacrylamide gel electrophoresis (PAGE) analysis **(A)**, and Quantification **(B)** of *Sel1L* exon 4 splicing in various mouse tissues. Statistics indicates the comparison between pancreas and other tissues. No statistically significant differences were observed between any of the other tissues. Hippo, Hippocampus. Cb, Cerebellum. BAT, Brown adipose tissue. n = 3-4 mice/group. **(C to D)** DNA polyacrylamide gel electrophoresis (PAGE) analysis **(C)**, and Quantification **(D)** of *Sel1L* exon 4 splicing in DMSO or Tunicamycin (Tuni)-treated mouse livers. n=3-4 mice/group. Data are represented as means ± SEM. n.s., not significant. *p < 0.05; **p < 0.01; ***p < 0.001, ****p < 0.0001. One-way ANOVA followed by Tukey's post hoc test for **(B),** two-tailed t test for **(D)**.

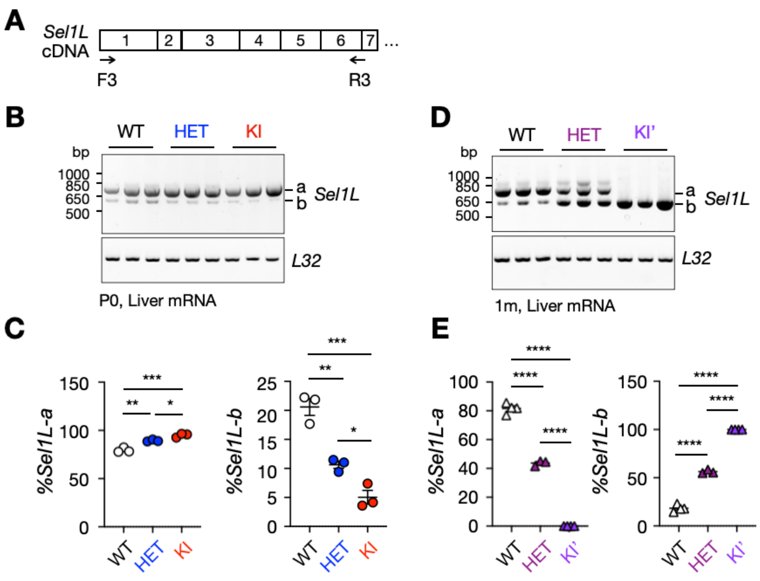


**Fig. S3. The *Sel1L* isoforms in KI and KI' mice. (A)** The diagram of the primer design for **(B)** and **(D)**. **(B to E)** The agarose electrophoresis **(B)** and **(D)** and Quantitation **(C)** and **(E)** of *Sel1L* isoforms in WT, HET, and KI/KI' mice. n = 3-4 mice/each group. Data are represented as means ± SEM. *p < 0.05; **p < 0.01; ***p < 0.001, ****p < 0.0001. One-way ANOVA followed by Tukey's post hoc test for **(C and E)**.

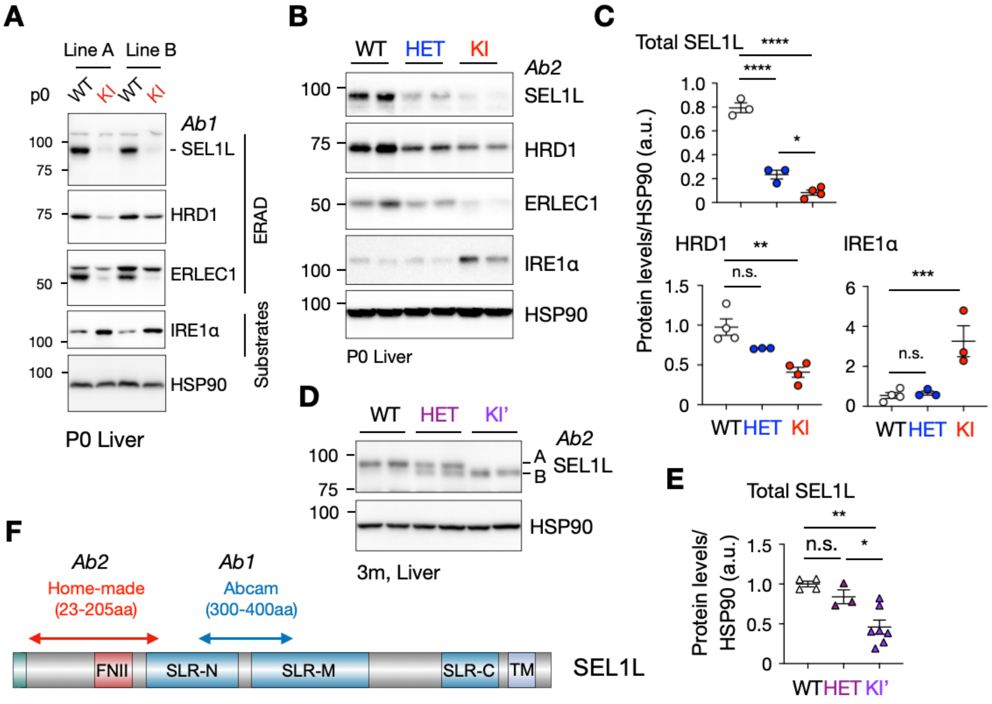


**Fig. S4. SEL1L C141Y KI mice showed ERAD deficiency in the liver. (A)** Western blot analysis of ERAD protein and ERAD substrates from WT and KI P0 pup liver from Line A and Line B. **(B and C)** Western blot analysis **(B)** and Quantitation **(C)** of ERAD protein and ERAD substrates from WT, HET, and KI P0 pup liver. n = 3-4 mice/group. **(D to E)** Western blot analysis **(D)** and Quantitation **(E)** of SEL1L from WT, HET, and KI' mouse liver. n=3-4 mice/group for KI line and 4-7 mice/group for KI' line. Sex and age combined. **(F)** Diagram showing the epitope target regions of the home-made and Abcam anti-SEL1L antibodies on the SEL1L protein. Data are represented as means ± SEM. n.s., not significant. *$p < 0.05$; **$p < 0.01$; ***$p < 0.001$; ****$p<0.0001$. One-way ANOVA followed by Tukey's post hoc test for **(C and E)**.

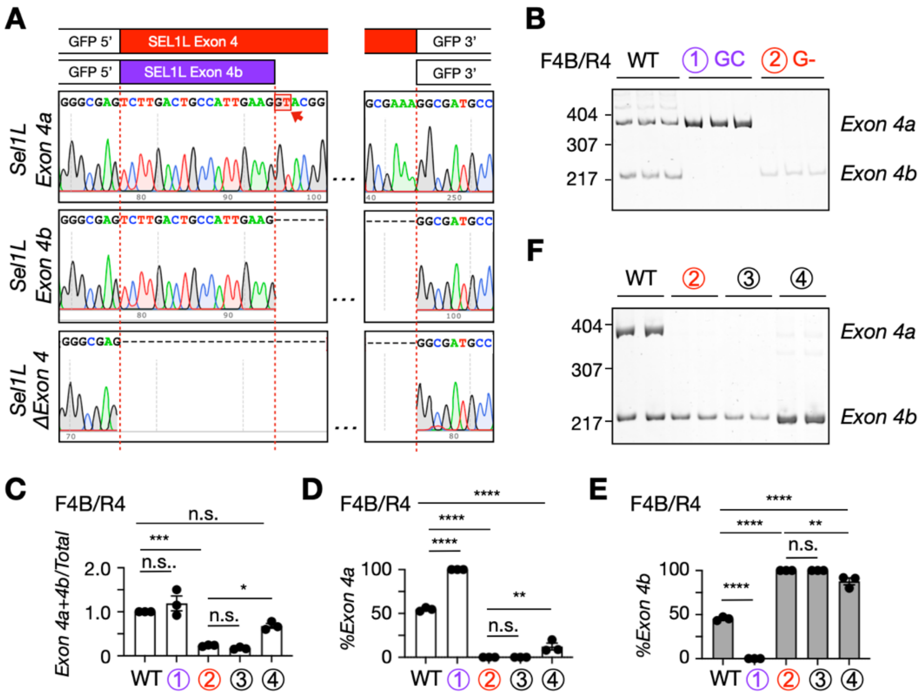


**Fig. S5. Confirmation of *Sel1L* exon 4 alternative splicing in the minigene reporter construct. (A)** Sanger sequencing confirmation of the splice products shown in **Fig. 2, E and H**. **(B and F)** DNA PAGE analysis of the WT and mutated minigene constructs as indicated by using primer pair F4B/R4. **(C)** Quantitation of *Sel1L* Exon 4a+4b transcript level in **(B)** and **(F)** normalized by total *Sel1L* transcript level as shown in **Fig. 2, E and H**. **(D and E)** Quantitation of **(B)** and **(F)** as percent of exon 4 inclusion. n = 3 independent biological replicates. Statistics indicates the comparison between WT and mutated constructs. $^{*}p < 0.05$; $^{**}p < 0.01$; $^{***}p < 0.001$, One-way ANOVA followed by Tukey's post hoc test for **(c),** Two-way ANOVA followed by Tukey's multiple-comparisons test for **(D to E)**.

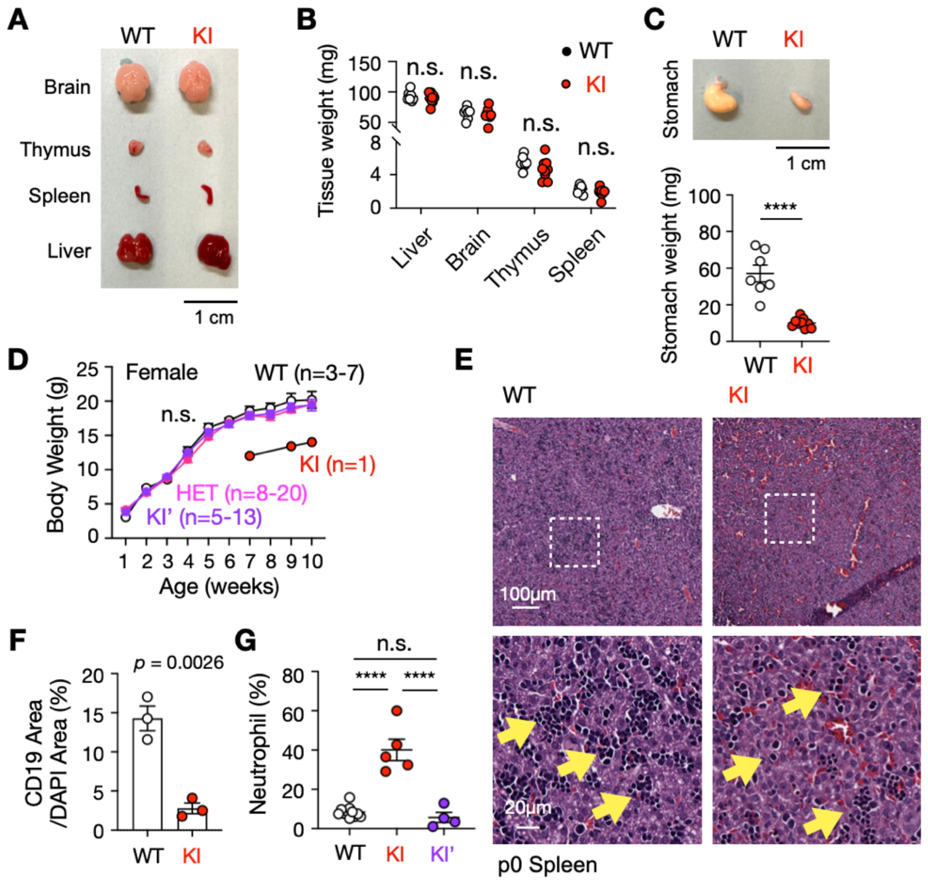


**Fig. S6. Tissue weight and B cell profile in the spleen of SEL1L C141Y KI mice. (A to B)** Tissue morphology **(A)** and tissue weights **(B)** of SEL1L C141Y knock-in (KI) pups at postnatal day 0 (P0). n = 7-9 mice/group. **(C)** Morphology and weight quantitation of postprandial stomach in the P0 WT and KI pup. n = 7-9 mice/group. **(D)** Body weight growth of female WT, HET, KI' and surviving KI mice for postnatal week 1 to week 10. **(E)** Hematoxylin and eosin (H&E) staining of the spleen from SEL1L C141Y KI pups at P0. Yellow arrow, lymphocytes. **(F)** Quantitation of CD19+ signal area normalized by DAPI + area of the immunofluorescence staining in **Fig. 3G**. n=3 mice/group. **(G)** Percentage of neutrophils in the peripheral blood mononuclear cells (PBMCs) of WT, KI' and surviving KI mice. Age and sex combined. n=4-11 mice/group. Data are represented as means ± SEM. n.s., not significant. ****p < 0.0001. One-way ANOVA followed by

Tukey's post hoc test for **(B and G),** two-tailed t test for **(C and F)**, Two-way ANOVA followed by Tukey's multiple-comparisons test for **(D)**.

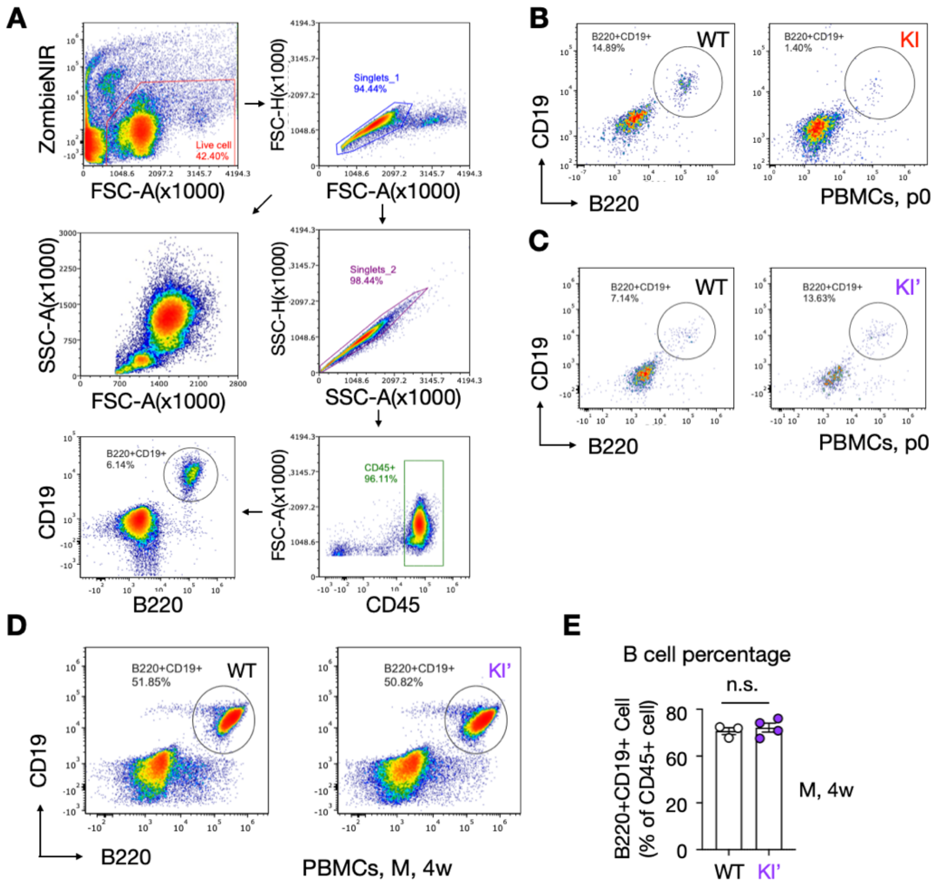


**Fig. S7. B cell deficiency in the circulation of SEL1L C141Y KI mice. (A)** Gating strategy for flow cytometry. Cells were first gated for live cells using Zombie NIR viability dye, followed by singlet gating to exclude doublets. Live singlet cells were then gated for CD45$^{+}$ leukocytes, and B cells were identified as CD19$^{+}$B220$^{+}$ within the CD45$^{+}$ population. **(B to D)** Representative flow cytometry plots showing CD19$^{+}$B220$^{+}$ B cells in PBMCs from WT and KI pups at postnatal day 0 (P0) **(B)**, and WT and KI' pups at P0 **(C)** and 4 weeks of age **(D)**. **(E)** Quantification of B cells as a percentage of CD45$^{+}$ cells as shown in **(D)**. n=3-4 mice/group. Data are represented as means ± SEM. n.s., not significant. Two-tailed t test for **(E)**.

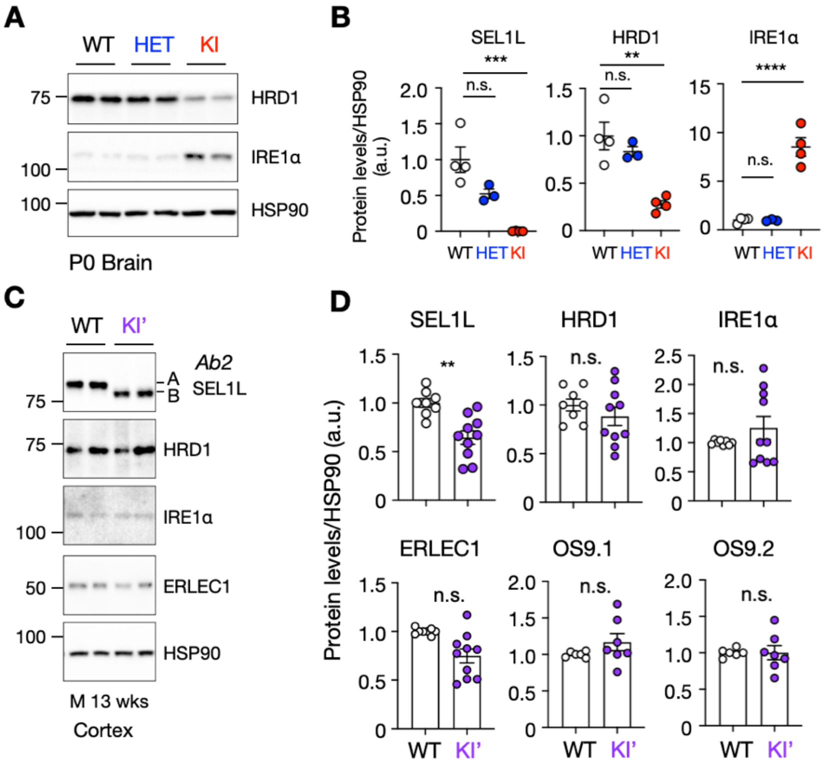


**Fig. S8. SEL1L C141Y KI mice showed ERAD deficiency in the brain, which was reversed in KI' mice. (A and B)** Western blot analysis **(A)** and Quantitation **(B)** of ERAD protein and ERAD substrates from WT, HET, and KI P0 pup brain with quantitation on the right. n = 3-4 mice/group. **(C and D)** Western blot analysis **(C)** and quantitation **(D)** of ERAD protein and ERAD substrates from the cortex of WT and KI' mice with quantitation on the right. n = 6-7 mice/group. Sex and age combined. Data are represented as means ± SEM. n.s., not significant. *p < 0.05; **p < 0.01; ***p < 0.001; ****p<0.0001. One-way ANOVA followed by Tukey's post hoc test for **(B)**, Two-tailed t test for **(D)**.

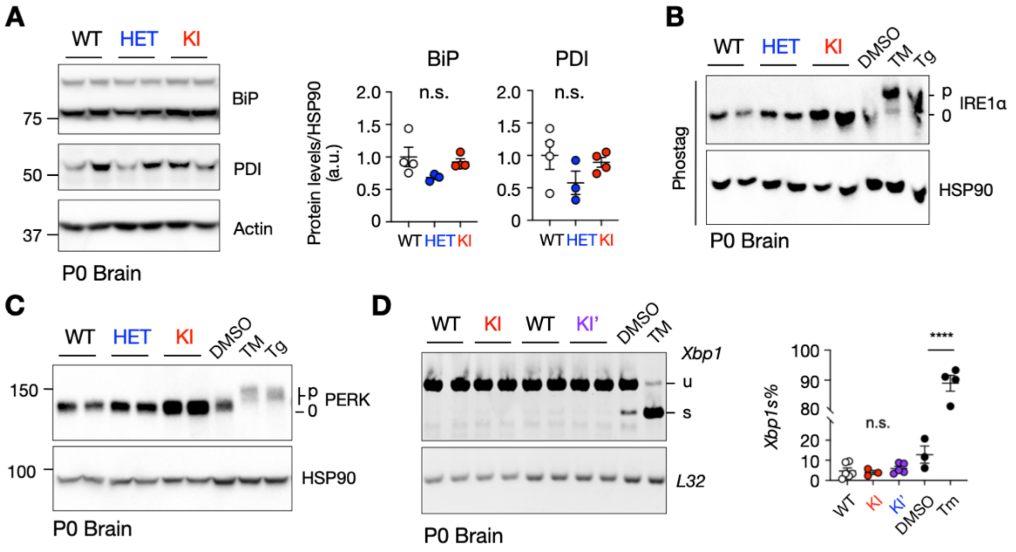


**Fig. S9. SEL1L C141Y KI mice did not cause overt UPR activation. (A)** Western blot analysis and quantitation of ER chaperones BiP and PDI in P0 WT, HET and KI mouse brains. n=3-4 mice/group. **(B)** Phos-tag gel analysis of IRE1α phosphorylation in P0 WT, HET and KI mouse brains. n=3-4 mice/group. **(C)** Western blot analysis of PERK in P0 WT, HET and KI mouse brains. n=3-4 mice/group. **(D)** Electrophoresis analysis of XBP1 splicing of P0 WT, KI, and KI' mouse brains with quantitation on the right. DNA PAGE gel for the *XBP1* splicing and agarose gel for the internal control *PPIA*. n=3-6 mice/group. n=3-4 for positive controls. p, phosphorylated. 0, un-phosphorylated. u, unspliced. s, spliced. Data are represented as means ± SEM. n.s., not signif-icant. ****p<0.0001. One-way ANOVA followed by Tukey's post hoc test for **(A)** and **(D)**.

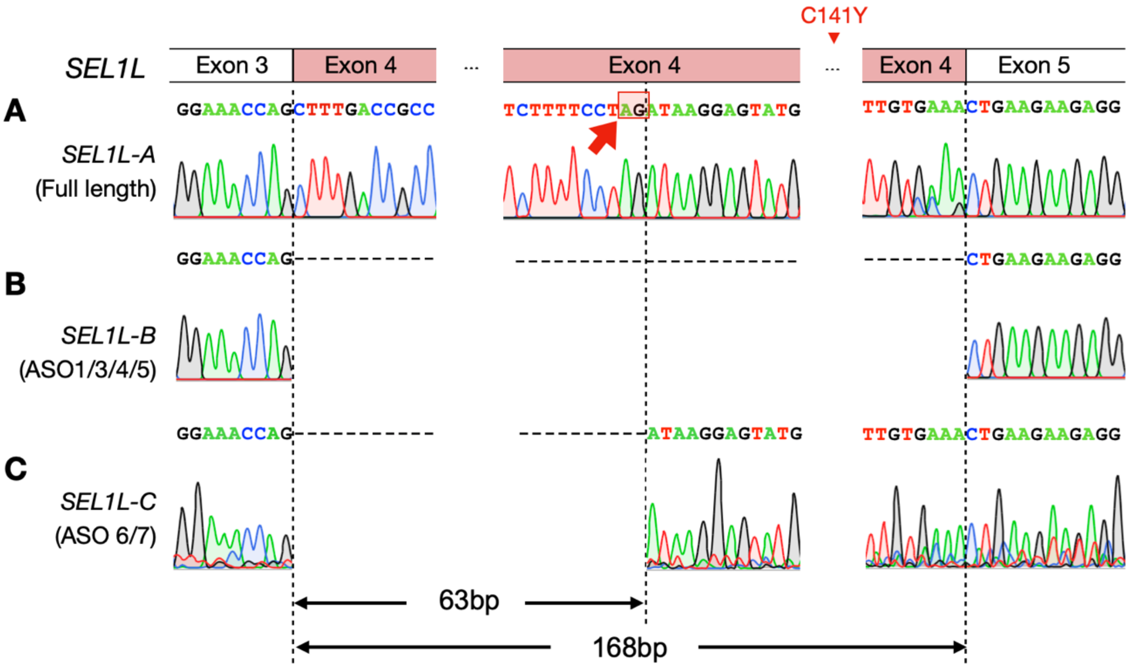


**Fig. S10. Sanger sequencing confirmation of ASO-mediated exon skipping in human patient fibroblast.** Sanger sequencing confirmation of full-length band or skipping bands from patient fibroblast treated with control oligo **(A)**, ASO1/3/4/5 **(B)**, and ASO6/7 **(C)**, as shown in **Fig. 4C**.

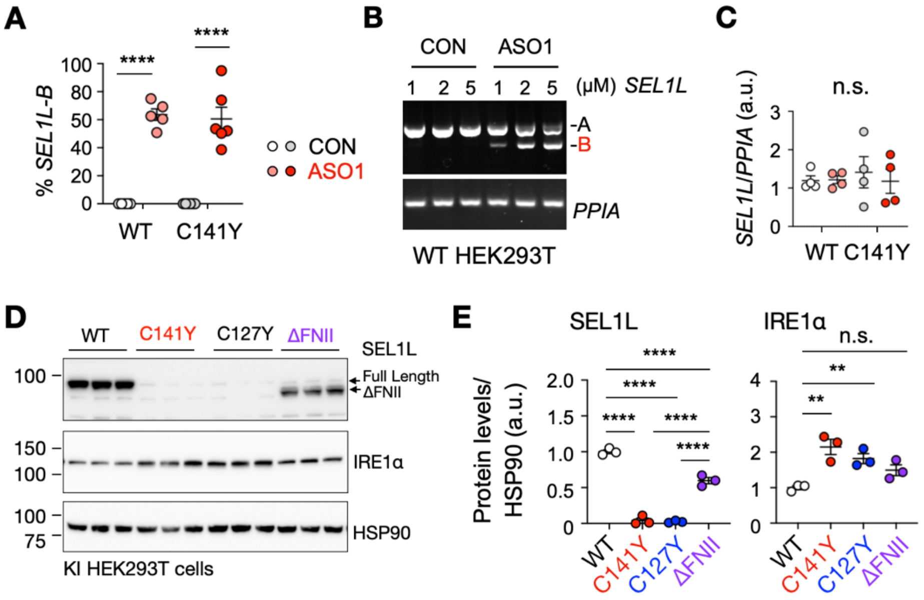


**Fig. S11. The regulation of *SEL1L* alternative splicing and exon skipping. (A)** Quantification of transcript isoforms from the agarose gel analysis shown in **Fig. 4E**. n = 5-6 independent biological replicates. **(B)** HEK293T cells were treated with control oligo or ASO1 at the indicated concentrations for 24 hours. SEL1L full-length and ΔExon 4 transcripts were analyzed by DNA agarose gel electrophoresis. **(C)** Quantitation of total *SEL1L* transcript level normalized by *PPIA* in **Fig. 4E**. n=4 independent replicates. **(D and E)** Western blot analysis **(D)** and Quantitation **(E)** of indicated proteins in WT, SEL1L C141Y, SEL1L C127Y and SEL1L FNII domain deletion (ΔFNII) HEK293T cells. n = 3 technical replicates. Data are represented as means ± SEM. n.s., not significant. *$p < 0.05$; **$p < 0.01$; ***$p < 0.001$, ****$p < 0.0001$. Two-way ANOVA followed by Tukey's multiple-comparisons test for **(A and C),** One-way ANOVA followed by Tukey's post hoc test for **(E)**.
